# EasyGrid: a versatile platform for automated cryo-EM sample preparation and quality control

**DOI:** 10.1101/2024.01.18.576170

**Authors:** Olivier Gemin, Victor Armijo, Michael Hons, Caroline Bissardon, Romain Linares, Matthew W. Bowler, Georg Wolff, Kirill Kovalev, Anastasiia Babenko, Veijo T. Salo, Sarah Schneider, Christopher Rossi, Léa Lecomte, Thibault Deckers, Kévin Lauzier, Robert Janocha, Franck Felisaz, Jérémy Sinoir, Wojciech Galej, Julia Mahamid, Christoph W. Müller, Sebastian Eustermann, Simone Mattei, Florent Cipriani, Gergely Papp

**Author notes:** contributed equally (Olivier Gemin, Victor Armijo).

## Abstract

Imaging biological macromolecules in their native state with single-particle cryo-electron microscopy (cryo-EM) or *in situ* cryo-electron tomography (cryo-ET) requires optimized approaches for the preparation and vitrification of biological samples. Here, we describe EasyGrid, a versatile technology enabling systematic, tailored and advanced sample preparation for cellular and structural biology. This automated, standalone platform combines in-line plasma treatment, microfluidic dispensing, blot-less sample spreading, jet-based vitrification and on-the-fly grid quality control using light interferometry to streamline cryo-EM sample optimization. With EasyGrid, we optimized grid preparation for different purified macromolecular complexes and subsequently determined their structure with cryo-EM. We also demonstrated how the platform allows better vitrification of large, mammalian cells compared to standard plunge-freezing. Automated sample preparation with EasyGrid establishes an advanced, high-throughput platform for both single-particle cryo-EM and cellular cryo-ET sample preparation.

## Introduction

Cryo-electron microscopy (cryo-EM) allows investigating the structure of macromolecular assemblies both in purified solutions using single-particle analysis^1^ (SPA) and inside cells using cryo-electron tomography^2-4^ (cryo-ET). Even with recent advances in cryo-EM/ET methods, the challenges associated with preparing high-quality vitreous samples remain a limiting factor.

Preparing an adequate sample for SPA — i.e. forming a thin film (< 50 nm) of vitreous water containing intact macromolecules and spread on a meshed support (grid) compatible with transmission electron microscopy (TEM) — requires the prior optimization of numerous parameters of sample preparation. Among these parameters, the thickness of the vitrified sample determines the signal-to-noise ratio (SNR) of TEM images. Sample thickness depends on the microfluidic behavior of the sample solution and on the wettability of the meshed support during grid preparation. Therefore, sample thickness is influenced not only by sample composition, viscosity and ionic strength but also by grid geometry, grid surface properties, sample handling and vitrification speed^5^. Due to the lack of data to model and control sample spreading on cryo-EM grids, it is common practice to screen the parameter space of sample preparation until producing adequate grids. New tools were developed to facilitate grid optimization: some instruments rely on the long-standing plunge-freezing approach^6^ while others harness microfluidic dispensing and/or ethane-jetting technologies^7-11^. Even with these tools, sample preparation remains a major bottleneck for most cryo-EM projects, emphasizing the need to develop automated screening of grid preparation parameters. Besides, the latest sample preparation instruments preclude vitrifying cells grown on grids for cryo-ET experiments — either because they can only handle grids that are already clipped in a copper cartridge^8^ which is toxic to cells, or because they are not equipped to remove culture medium from the cell-seeded grid before vitrification^7,9-11^. Additionally, plunge-freezing often fails to fully vitrify thick cellular samples due to insufficient cooling rates inside cells resulting in sample crystallization deteriorating cellular ultrastructure^12^. In contrast, actively jetting cryogen on non-clipped grids after quickly removing liquids from cell-seeded grids would yield higher cooling rates allowing to vitrify bulkier samples^13,14^. In summary, a modular and automated framework entailing all steps of grid production and including ethane-jetting technology is required to streamline grid optimization.

Beyond sample preparation, grid quality control is a major bottleneck of sample optimization because it still requires expensive access to cryo-EM instrumentation. In more detail, the standard procedure for controlling the quality of SPA grids is a two-step imaging process. An overview map (atlas) of the grid is first acquired at ∼100x magnification (∼120 nm/pixel) to identify grid regions with adequately low sample thickness. Grids covered with thick or crystalline ice are readily discarded to make time for valuable grids. Secondly, a ‘high-resolution’ quality control is performed by imaging a selection of grid squares at typically >40,000x magnification (<0.5 nm/pixel) to assess particle concentration, distribution and homogeneity, as well as potential ice contamination. Notably, only the first of these steps (sample thickness mapping) is necessary to control the quality of cell-seeded cryo-ET grids. Therefore, an instrument capable of performing ice thickness mapping at cryo-temperature right after grid production would considerably speed up sample optimization for all cryo-EM pipelines. However, cryo-light microscopes (cryo-LM) lack the axial resolution necessary to characterize sample thicknesses below 400 nm because of the diffraction of light. Yet, digital holographic microscopy (DHM) is a light-based interferometry method that can map sample thickness variations in the nm-to-µm range^15^. Adapting DHM to cryogenic temperatures would enable inexpensive grid quality control before requesting cryo-EM beamtime.

To address the challenges of cryo-EM/ET sample preparation, we developed EasyGrid, a fully automated and modular platform for high-throughput sample vitrification using ethane-jetting technology. EasyGrid is designed to provide fine control over sample preparation parameters and to perform automated grid quality control at cryogenic temperature. We demonstrated that it is suited for optimizing SPA grids by obtaining high resolution maps of three different macromolecular complexes (i.e. apoferritin, yeast ribosomes and pentamers of KR2 bacterial rhodopsin). We further validated that large human cells exhibited better vitrification quality when prepared with EasyGrid compared to plunge-freezing. Thus, EasyGrid provides new solutions to improve and accelerate cryo-EM studies both *in vitro* and *in situ*.

## Results

### EasyGrid modules automate sample preparation and quality control

EasyGrid is a standalone platform equipped with multiple modules that perform all key functions for cryo-EM sample preparation, vitrification and short-term storage at liquid nitrogen (LN_2_) temperature (Suppl. Movie S1). EasyGrid includes the following modules:

#### Grid loading and handling module

Sample supports are first loaded in the machine using a custom grid rack (Fig. 1a, first panel). Compatible supports entail 3 mm-diameter cryo-EM grids covered with a perforated support foil. Individual grids are picked up by the automated grid handling module composed of a metallic gripper jaw motorized by a three-axis cartesian robot that drives the grid to the other modules. A system of two cameras set up perpendicularly to the gripper axis is used to calibrate gripper position and to monitor grid gripping before and during grid preparation. Automated grid handling reduces risks of grid bending and the deterioration of the support foil.

**Figure 1.**
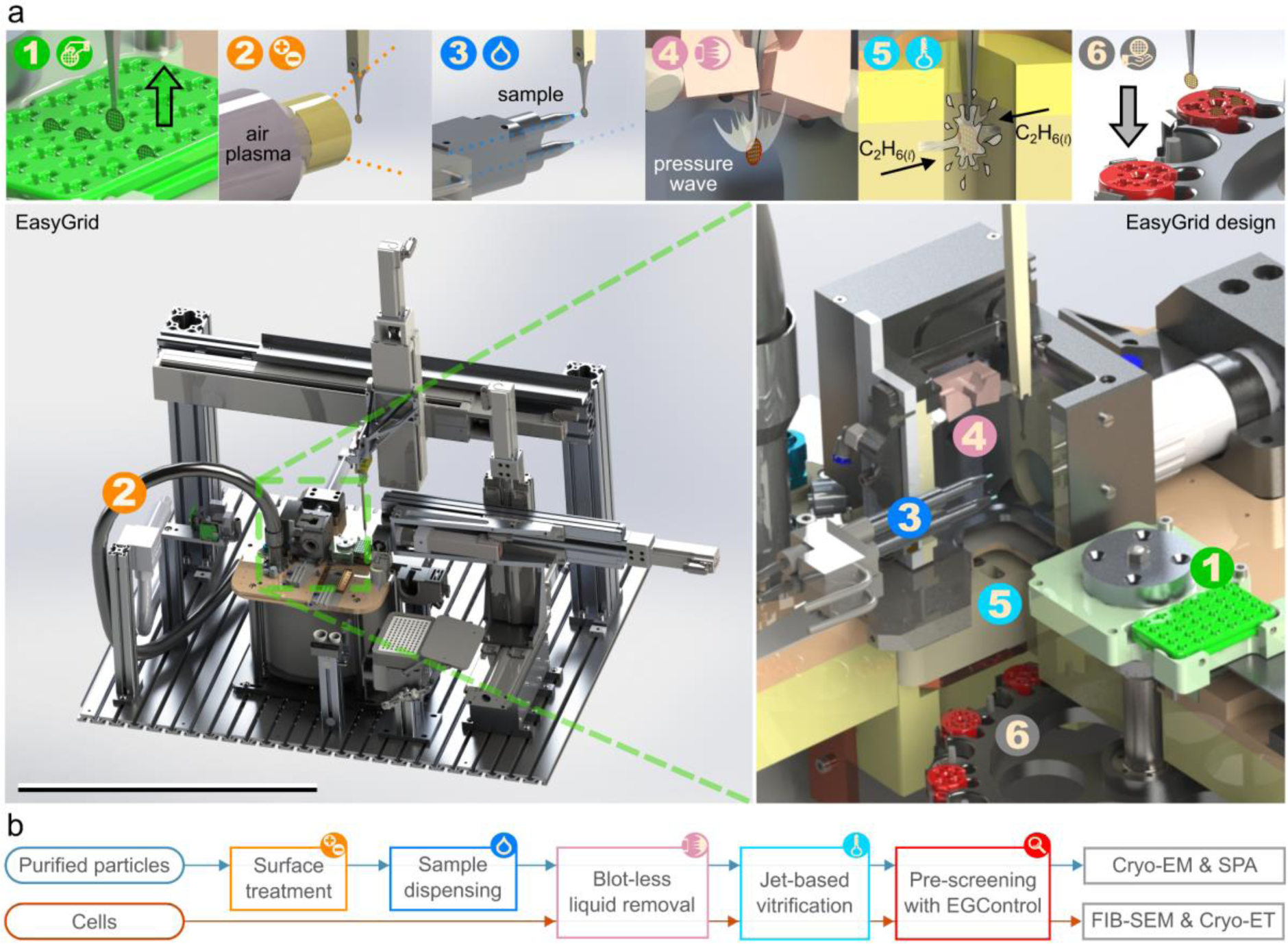
EasyGrid, a modular platform combining key technologies for cryo-EM grid preparation and quality control. a. Overview of the EasyGrid platform. Upper row: schematics of EasyGrid modules. 1: Grid loading rack. 2: Plasma torch. 3: Two-nozzled microfluidic dispenser. 4: Pressure wave generator. 5: Ethane-jetting slot. 6: Grid-storing carousel. Lower row: rendering of the EasyGrid machine and close-up on the high-humidity chamber. Live close-ups in Suppl. Movie S1; scale bar, 1 m. b. Flow-chart of the two main workflows developed for EasyGrid, which aim at vitrifying purified protein solutions for cryo-EM and single-particle analysis (SPA) or vitrifying cells for *in situ* imaging such as cryo-tomography (cryo-ET) of lamellae milled with a focus ion beam/scanning electron microscope (FIB-SEM).

#### Plasma treatment module

EasyGrid is equipped with an atmospheric plasma treatment module (Fig. 1a, second panel) enabling the rapid and controlled enhancement of grid hydrophilicity. The water-cooled plasma torch is set up on a fixed holder, oriented perpendicularly to the grid and it outputs a power of 1 kVA at a frequency of 21kHz. During plasma treatment, the grid is moved back and forth several times 12 mm in front of the torch to charge the surface of the grid without damaging the gripper. Fifteen such passages (∼10 s total duration) are typically performed before delivering sample solutions on the grid.

#### Dual-nozzle picoliter drop dispenser

Sample solutions are provided by the user in a 96-wells PCR-format plate inserted in a covered and thermo-regulated dock (4 °C - 36 °C). Sample pipetting and deposition on the grid is performed automatically by the pipette robot (Fig. 1a, third panel) composed of two custom-designed, motorized and temperature-controlled (4 °C - 36 °C) piezo-electric picoliter drop dispenser heads. Once charged with >15µL of solution, the piezo-electric elements of the dispenser heads are first tuned to achieve monodisperse droplet dispensing using a stroboscopic observation module synchronized with drop dispensing events. The dispenser heads can then deliver droplets of ∼50 pL at a rate of up to 1000 droplets per second at custom locations on the grid. At least 30 grids can be prepared for a single charge of sample solution.

#### Humidity-controlled, pressure-wave-based sample spreading module

Direct freezing of sample droplets dispensed on grids resulted in samples that were too thick for TEM imaging. To prepare thin sample layers, EasyGrid is equipped with a pressure wave generator composed of two nozzles aiming at each side of the grid from a grazing angle (Fig. 1a, fourth panel). These nozzles are mounted in a small temperature-regulated chamber (4 °C - 36 °C) separated from the vitrification module by a trapdoor. High humidity (typically >85%) is maintained in this chamber to limit sample evaporation. The nozzles conveying the pressure wave to the grid are powered with oil-free, filtered, compressed air and operated with an over-sampled digital output module (∼1 µs switching time) ensuring the programmability and repeatability of pressure pulses. Sample spreading is performed by positioning the grid at the aiming point of the pressure wave generator and releasing a brief blast of air to sweep both sides of the grid. Typical spreading durations are 30 ms for biochemical solutions and 300 ms to remove culture medium from a cell-seeded grid. After spreading and an optional, adjustable time delay, the grid is driven within 200 ms to the ethane-jetting vitrification module.

#### Vitrification and storage module

The sample vitrification and storage module is composed of a dewar that is automatically topped-up with LN_2_, a temperature-regulated ethane reservoir with two output orifices facing each other (Fig. 1a, fifth panel), an ethane recycling pool and a grid storage carousel (Fig. 1a, sixth panel). Ethane is liquefied in the reservoir by flowing ethane gas through it. A single charge of ethane lasts for a whole day of operation thanks to the ethane recycling device. Ethane jets are generated by pressurizing the ethane reservoir to push cryogen towards the jetting orifices. Once colliding jets are stable for ∼500 ms, the grid is driven to the jets’ collision point to achieve sample vitrification. The grid is then kept in nitrogen vapor (-180 °C) to prevent devitrification and to limit ice contamination. The grid is finally inserted in a slotted box of the storage module, composed of a carousel that can accommodate up to 10 EasyGrid cryo-boxes. These custom boxes contain four grid slots and a cryo-compatible RFID chip (eCryoID^TM^ tag) that facilitates sample tracking and grid inventory.

The modules described above can be programmed into custom workflows to prepare two types of samples: solutions of purified particles for SPA, or cells for *in situ* cryo-ET (Fig. 1b).

#### Grid pre-screening with the EasyGrid Control module

A first criterion of grid usability for SPA is an appropriate thickness of the vitrified sample, which can be assessed with the EasyGrid Control (EGC) module. EGC is either integrated in the EasyGrid platform or built as a standalone ice thickness mapping device. In this case, the motorized cryo-observation column is installed on top of a carousel similar to that of the EasyGrid main machine (Fig. 2a, left panel) with a capacity for 40 grids that are sequentially picked up by a motorized gripper (Fig. 2a, middle panels). To maintain the gripper temperature stable below -170 °C, the column inner tube is made of copper, immersed in LN_2_ and thermally insulated from the exterior. Portholes of high optical quality installed across the observation point (Fig. 2a, right panel) enable mapping sample thickness with the interferometry setup depicted in Fig. 2b: a laser beam is split into an unmodulated “reference” and a “measurement” beam that illuminates the grid. The variations of refractive index and sample thickness across the grid modulate the wavefront of the measurement beam. The two beams are re-combined to interfere with each other, and the resulting hologram is recorded on a CCD camera. Unwrapping and analyzing the phase of the hologram allows deriving sub-wavelength measurements of optical thickness across the grid. Currently, EGC comprises three complementary imaging modes:

**Figure 2.**
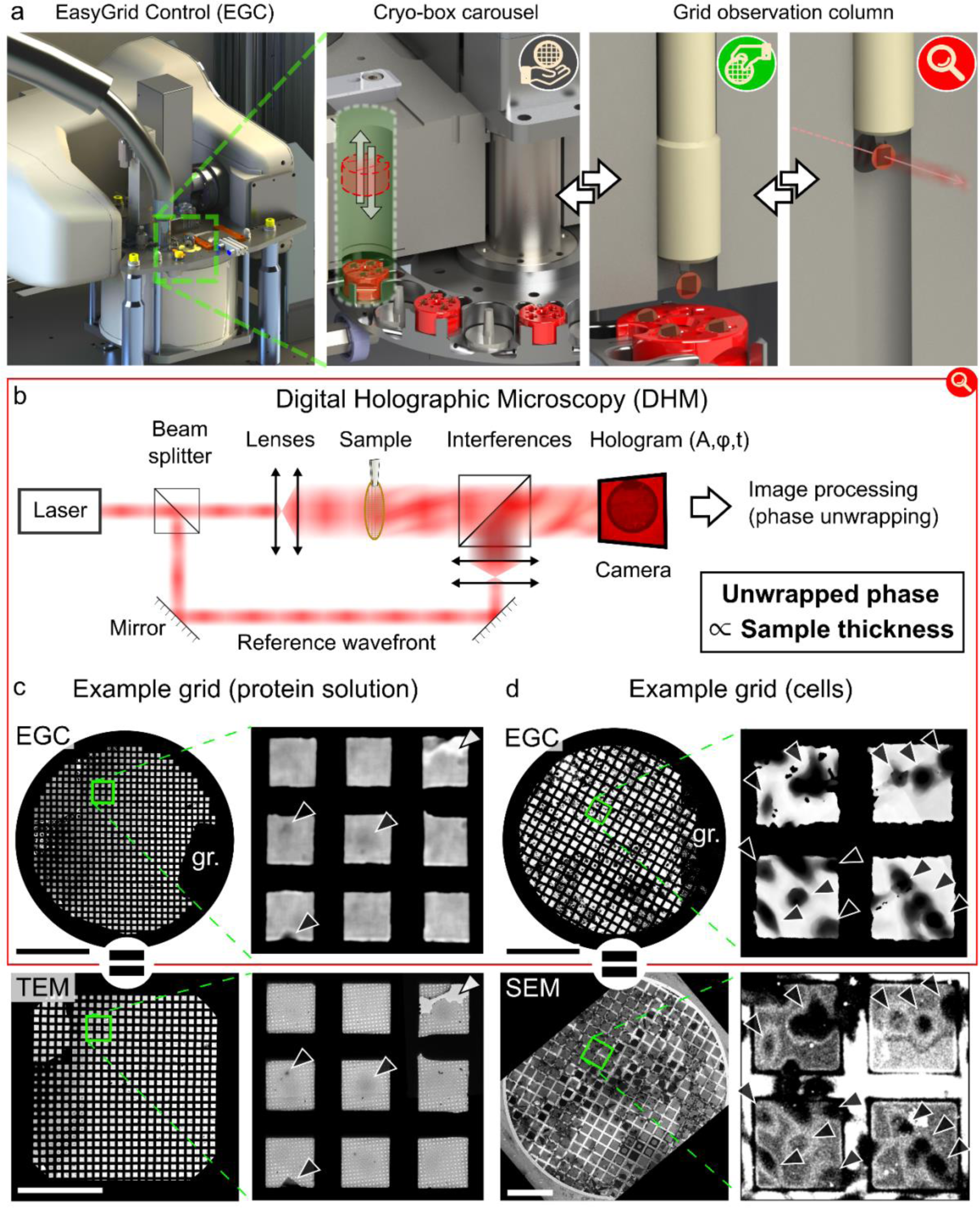
EasyGrid Control enables ice-thickness mapping at cryo-temperature to determine optimal grids for cryo-EM imaging. a. Schematics of the EasyGrid Control module (first panel to the left) and close-ups on: the grid box exchange trapdoor (highlighted in green with white arrows representing box exchange by the user) and cryo-carousel (2^nd^ panel), the grid pick-up point (3^rd^ panel) and the grid observation point (4^th^ panel). b. Principle of digital holographic microscopy to acquire thickness maps with sub-wavelength axial resolution using interferometry. c. Atlases and close-ups of one grid prepared for single-particle analysis and imaged with both EGC (top panels) and cryo-EM (bottom panels). Insets show features visible in both modalities. White arrowhead: a hole in the carbon film. Black arrowheads: local maxima of ice thickness. More examples in Suppl. Fig. S1; gr: grid gripper; scale bars, 1 mm. d. Atlas of one cell-seeded grid imaged with both EGC (top) and cryo-SEM (bottom). The insets show cells (arrowheads) visible in both modalities. More examples in Suppl. Fig. S1; gr: grid gripper; scale bars, 1 mm (top) and 500 µm (bottom).

##### Thickness mapping at 2x magnification (∼3 µm/pixel)

This observation mode allows rapidly (∼5’/grid) assessing overall sample quality and ice contamination.

##### Screening at 10x magnification (∼0.6 µm/pixel)

Because of the limited field of view at 10x magnification (∼0.35 mm x 0.35 mm), grids are imaged as a montage of 9x9 tiles (∼15’/grid).

##### Thickness mapping at 10x magnification

Deriving thickness measurements from a 10x tile requires a thickness reference for each tile (e.g. “0 nm” measured in a broken square) but such references are difficult to propagate across the montage because of measurement discontinuities (e.g. grid bars or thick squares). We addressed this issue by calibrating the average thickness value of each 10x tile using a 2x map of the same grid. Calibrated 10x maps reliably correlate with cryo-EM atlases and describe the distribution of ice thickness at the level of individual squares for both SPA and cryo-ET samples (Fig. 2c-d). Thus, EGC allows rapid, inexpensive and automatic screening of batches of up to 40 grids to bring forward optimal specimens (Suppl. Fig. S1) for subsequent cryo-EM/ET imaging.

### EasyGrid provides high-quality samples for structural biology

#### EasyGrid automates cryo-EM grid preparation for single-particle analysis

The aim of SPA workflows is to derive the 3D structure of molecular assemblies from thousands of cryo-EM images of macromolecular complexes immobilized in a thin vitrified ice layer. Preparing large areas of sample film that are suitable for high-resolution cryo-EM is challenging in practice: exploratory optimization is often required to simultaneously obtain a minimal thickness of the frozen sample layer, a dense particle distribution, varied particle orientations, and the absence of ice crystals. To facilitate SPA sample optimization, we developed a versatile and reproducible workflow encompassing all steps of grid production (Fig. 3a). The EasyGrid SPA routine starts with enhancing the hydrophilicity of a grid using atmospheric plasma treatment, then dispensing the sample or a pre-coating solution on the grid in a user-designed pattern (typically a line of 100 droplets) using microfluidic nozzles (Fig. 3b). A rapid sequence of sample spreading with the pressure-wave system followed by jet-vitrification allows freezing unstably thin sample films^16^ on grids (Fig.3b, right panel) before storing them in cryo-boxes at LN_2_ temperature. We implemented SPA grid preparation protocols to study human apoferritin, yeast ribosomes, and bacterial KR2 rhodopsin samples (Fig. 3c-e) and determined their structures at 1.9 Å, 2.4 Å, and 2.3 Å, respectively using conventional cryo-EM acquisition and processing schemes (Suppl. Figs. S2, S3 and S4). The structure of the KR2 rhodopsin complex was not previously described with cryo-EM although a structure obtained at high salt concentrations with X-ray crystallography was already available^17^. Our cryo-EM map at 2.3Å resolution shows key features of KR2 rhodopsin including the interprotomeric sodium binding site and the functional, expanded conformation of the central region of the pentamer with numerous water molecules populating the internal cavities. The cryo-EM map additionally reveals all 4 of the water molecules located in the Schiff base cavity of KR2 while only 3 water molecules were resolved in X-ray crystallography data at a comparable map resolution (2.2Å)^17^. Last, we observed a different organization of the hydrophobic tails of lipid molecules in the interprotomeric clefts compared to published crystal structures, suggesting the high flexibility of lipid-protein interactions in KR2 complexes (Suppl. Fig. S5).

**Figure 3.**
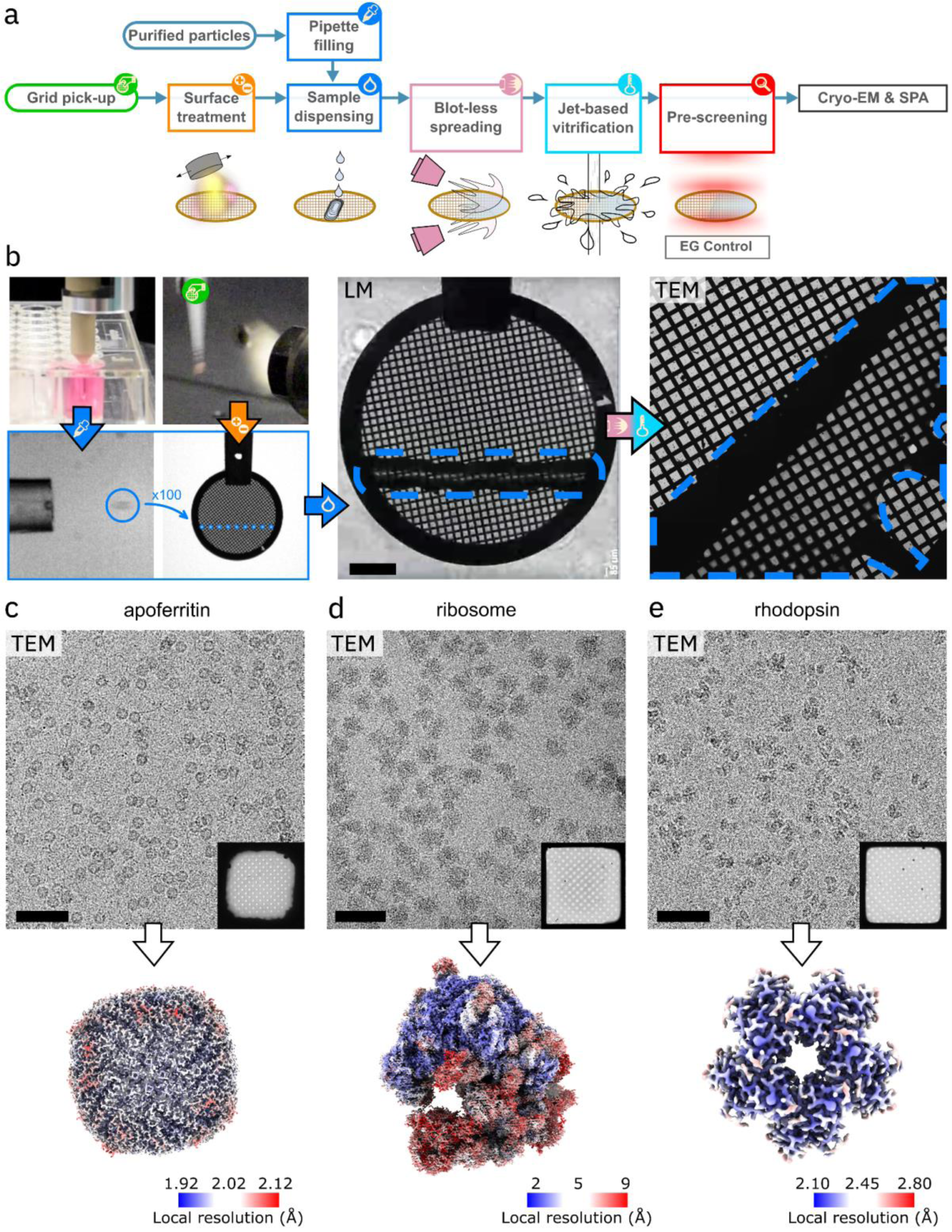
Automated grid preparation with EasyGrid enables high-quality cryo-EM acquisition and structure determination. a. Flow-chart of SPA grid preparation with EasyGrid which entails conditioning grids for optimal sample spreading before dispensing, spreading and jet-vitrifying a sample solution on the grid for subsequent cryo-EM imaging. Optional on-the-fly grid screening with EasyGrid Control (EGC) allows grid quality control prior to cryo-EM imaging. b. Snapshots of a SPA grid preparation. Initial steps include 1) filling the pipette robot with the sample solution (top-left panel) then tuning the piezo-electric dispensing system to reproducibly dispense sample droplets (blue circle) and 2) treating the picked-up grid with plasma (2^nd^ panel) and optional pre-coating steps (not shown). The pipette robot is then programmed to dispense droplets at desired locations (small cyan circles), typically to form a line of ∼10 nL of solution close to the midline of the grid (dashed blue line in the middle panel; scale bar, 500 µm). After sample spreading and vitrification, the grid can be imaged in cryo-EM (right panel; the dashed blue line outlines the region where the dispensed sample was spreaded). c-e. Top panels: Example micrographs acquired on grids prepared with EasyGrid using apoferritin (c), yeast ribosomes (d) or KR2 rhodopsin (e) solutions. Insets show grid squares in which the high-resolution micrographs were acquired; scale bars, 50 nm. Bottom panels: cryo-EM density maps color-coded by local resolution, reconstructed after performing SPA on grids prepared with EasyGrid.

In summary, we showed that EasyGrid can prepare high-quality SPA grids, allowing discovering macromolecular structures through fully automated processes that decrease human error factors in grid preparation efforts.

#### EasyGrid improves cell vitrification quality for *in situ* cryo-ET

Sample vitrification without crystalline ice formation is required to preserve the integrity of nanometric, subcellular features for high-quality cryo-ET imaging, but plunge-freezing provides insufficient cooling rates to fully vitrify samples thicker than a few µm. Jet-vitrification, as implemented in EasyGrid (Fig. 4a), was proposed to improve cooling rates^8,13,14^. To demonstrate the applicability of our cryo-ET sample preparation workflow, we cultured HeLa cells on micropatterned grids made of a gold mesh overlaid with perforated SiO_2_ support film, vitrified them with EasyGrid and then used a dual-beam focused ion beam/scanning electron microscope (FIBSEM) to produce thin lamellae following established protocols^18-20^ (cryoFIB-milling; Fig. 4b). Cryo-EM overviews of the lamellae and the subsequent tilt-series acquisitions exhibited no crystalline ice artifacts in the central regions of cells, which often displayed crystalline artifacts in lamellae milled from plunge-frozen cells (Suppl. Fig. S6). The tomograms we reconstructed from these cells did not exhibit crystalline artifacts (Fig. 4c).

**Figure 4.**
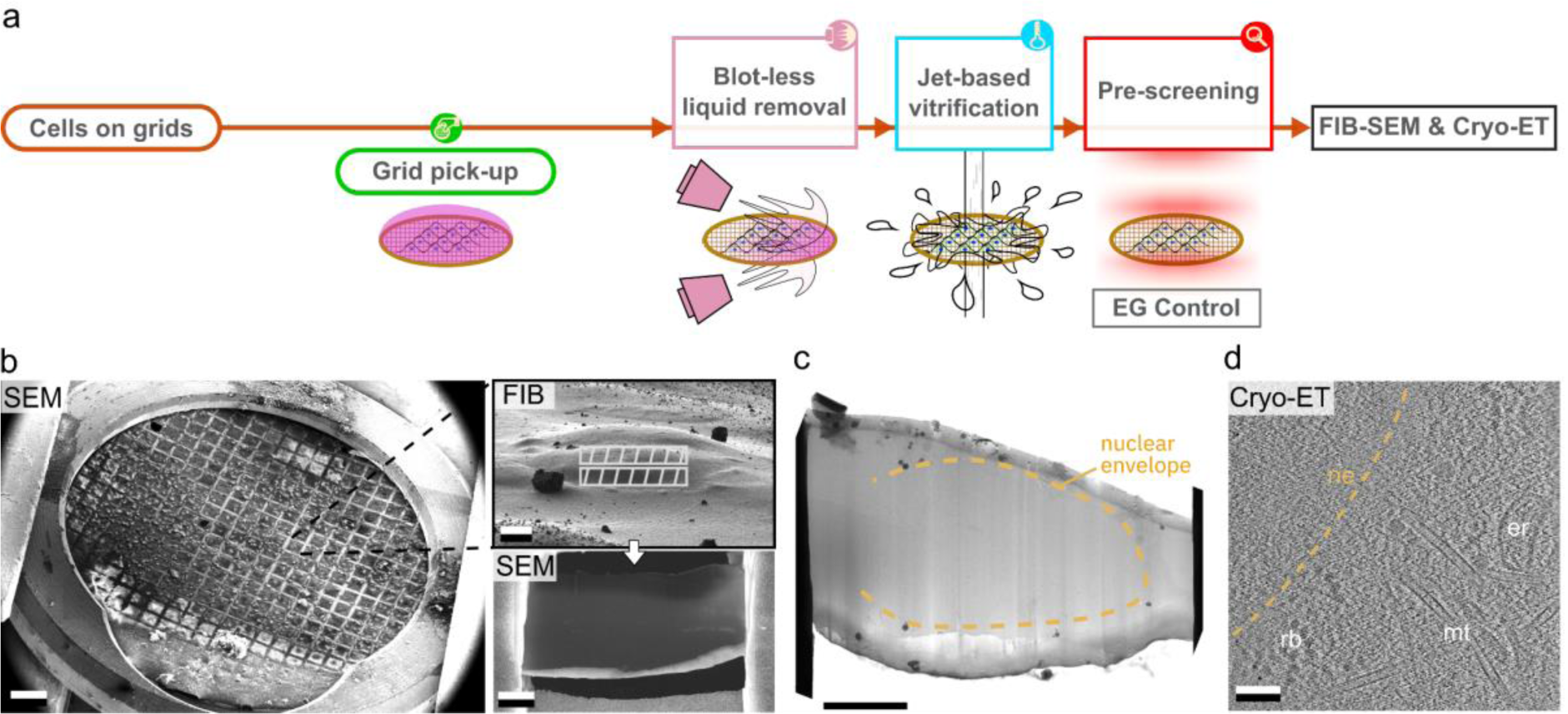
EasyGrid provides high quality samples for *in situ* cryo-ET in large cells. a. Flow-chart of cell-seeded grid preparation from growing adherent cells on cryo-EM grids to sample thinning, vitrification, and quality control, followed by FIB lamella preparation and cryo-ET. b. Example of lamella preparation in HeLa cells that were jet-vitrified on a cryo-EM grid and observed using SEM (scanning EM; left panel). The focused ion beam (FIB) image (top panel) is tilted 52° from the SEM optical path and shows the areas that were FIB-milled (gray patterns) to produce an electron-transparent lamella (lower panel) in the bulk of the cell; scale bars, 300 µm (left), 3 µm (top) and 2 µm (bottom). c. TEM overview of a lamella milled in the bulk of a HeLa cell prepared with EasyGrid. The absence of crystalline artifacts on the lamella illustrates good ice quality at a mesoscopic level; scale bar, 5 µm. d. Central section of a cryo-tomographic reconstruction of a HeLa cell prepared with EasyGrid; scale bar, 100 nm; er: endoplasmic reticulum; ne: nuclear envelope; mt: microtubule; rb: ribosomes.

To characterize the improvement of ice quality in cells prepared with EasyGrid compared to plunge-freezing, we first conducted a qualitative study on mammalian SUM159 cells (Suppl.Fig. S7) and performed a quantitative analysis of lamellae FIB-milled at mid-height of HeLa cells. Low magnification overviews of most lamellae milled in plunge-frozen cells displayed crystalline ice artifacts, whereas the majority of lamellae milled in jet-vitrified cells appeared vitreous (Fig. 5a and Suppl. Fig. S6). To quantify crystallinity, we carried out systematic cryo-EM raster imaging of the milled lamellae at a pixel size of 1.2Å and analyzed the 2D Fourier transform of these images to visualize the diffraction of ice around 0.27Å^-1^ (often described as the “ice ring at 3.7Å”)^21^. We excluded regions contaminated with large exogenous ice particles to restrict our analysis to electron-transparent regions of the lamellae (Fig. 5a, lower row). Performing peak detection analysis in the water ring region of the 2D power spectra of acquired images allowed categorizing each of the 5968 sampled positions as either “unambiguously vitreous”, “semi-crystalline” or “unambiguously crystalline” (Suppl. Fig S8). Our analysis showed that 77±14% of positions were vitreous in cells prepared with EasyGrid (N=16 lamellae) compared to 11±24% in cells prepared with standard plunge-freezing (N=20 lamellae; Fig. 5b). These results indicate that cells vitrified with EasyGrid displayed improved ice quality compared to plunge-frozen preparations, paving the way to further ultrastructural characterization of large cellular specimens.

**Figure 5.**
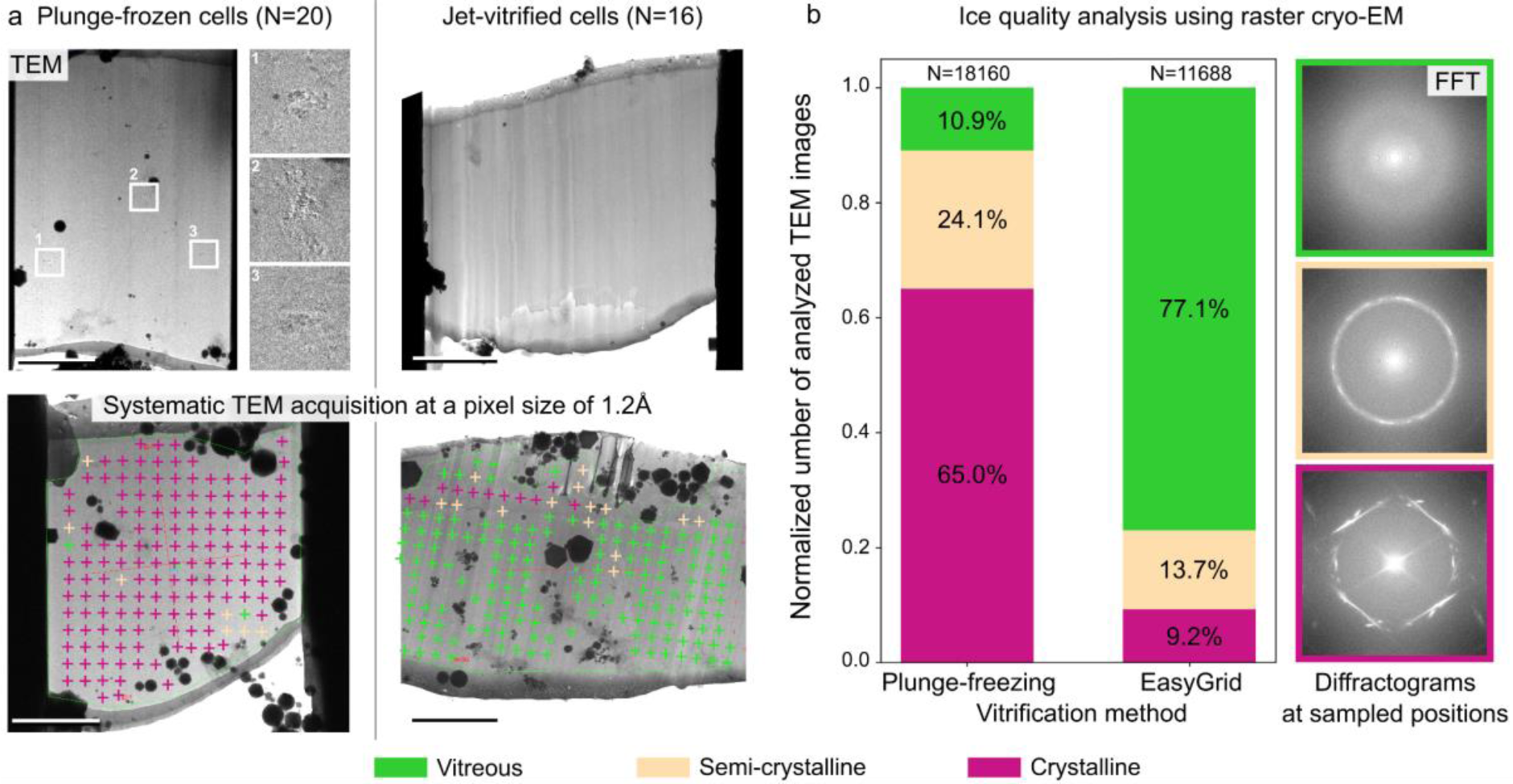
Jet-vitrification improves the vitreous quality of ice in lamellae milled in large mammalian cells. a. Cryo-EM maps of lamellae milled in the bulk of either plunge-frozen or jet-vitrified HeLa cells (left and right columns respectively). Insets in the upper left panel highlight crystalline ice artifacts that are already visible at 2250x magnification (5.6 nm/pixel). Crosses in the lower row represent positions where cryo-EM images were systematically acquired at 105,000x magnification (0.12 nm/pixel) to quantify ice quality across lamellae. The color code added *a posteriori* is “unambiguously vitreous”: green, “semi-crystalline”: orange, “unambiguously crystalline”: magenta; scale bars, 5 µm. b. Quantitation with EM diffraction analysis of ice crystallinity in lamellae prepared by either plunge-frozen or jet-freezing from Tokyo HeLa cells (N=20 and 16 lamellae, respectively). Insets show 2D Fourier transforms of images acquired in three different positions and displaying either a faint water ring without peaks in vitreous regions (top inset); stronger signal in the water ring with possible peaks in semi-crystalline and/or thicker regions (middle inset); or strong peaks at the spatial frequencies of water in crystalline regions (bottom inset with a hexagonal ice diffraction pattern indicating low local cooling rate^12^). No p-value was calculated for this categorical data.

## Discussion

Here, we presented EasyGrid, a modular technology that addresses major challenges of cryo-EM sample preparation and enables systematic, high-throughput cryo-EM/ET. We first demonstrated how EasyGrid allows optimizing macromolecular samples for SPA and subsequently determined three structures at about 2 Å resolution. We then showed how EasyGrid improves the vitrification of large cells, thus increasing the throughput of cryo-ET workflows. The versatility of EasyGrid and the ability to automatically assess sample quality with EGC makes this technology stand out from other grid preparation instruments.

Sample spreading in EasyGrid differs from blotting, wicking and pin-printing. Blotting is slow (seconds) and it possibly releases contaminants on grids^22^. By contrast, our pressure-wave system allows spreading samples within tens to hundreds of milliseconds and does not leach particles in spread samples. Secondly, wicking and pin-printing are not suited to prepare large adherent cells for cryo-ET but we demonstrated that EasyGrid is. Last, SPA grids prepared with self-wicking grids seemed to exhibit lower particle concentration than grids prepared with the Vitrobot^23^. By contrast, we did not observe concentration differences between grids prepared with EasyGrid or with the Vitrobot using the selfsame sample solution. EasyGrid might therefore also benefit from a concentration effect whereby particles accumulate in grid holes during sample preparation. Moreover, pressure-wave-based spreading did not prevent us from solving macromolecular structures at ∼2Å resolution, suggesting that EasyGrid is compatible with a majority of samples.

EasyGrid robotics ensure precise grid positioning, accurate timing of operations and consistent repeatability, which minimizes variability at each step of sample preparation compared to semi-automated preparation. Precisely controlling SPA sample delivery (±50 pL) and time-critical steps such as the delay between spreading and vitrification (minimum 200 ms in the current implementation; can be increased with a precision of ±1 ms) allows fine-tuning sample thickness using tailored dispensing and/or controlled evaporation of the specimen on the grid before freezing. However, chaotic phenomena (e.g. the microfluidic behavior of SPA samples, the seeding of cells on cryo-ET grids and the nucleation of ice crystals) singularize each vitrified sample. We propose automated sample preparation and quality control to overcome the irreducible variability of cryo-EM samples.

EasyGrid Control provides grid atlases enabling sample screening. The measurement of absolute sample thickness across grid squares is however still imprecise because of imaging artifacts (light diffraction by grid bars and the support film) and possible variations of optical properties (refractive index, support foil thickness) affecting acquired holograms. We are currently implementing higher magnification optics to alleviate artifacts and pursue ice quantitation. As of now, EGC assists in discarding subpar samples and in identifying high-quality grid areas. Once paired up with a neural network trained at analyzing atlases, EGC will become instrumental to fully automate grid optimization.

To approach samples that are currently difficult to prepare for SPA, we will upgrade EasyGrid to perform automated screens of biochemical parameters — i.e. optimize samples of incrementally varying composition following the design of protein crystallization platforms. One could combine this functionality with sample mixing^24^ to generate libraries of samples in which target macromolecules interact with arrays of selected ligands e.g. to develop new drugs. Further progress includes the insertion of a light injection line in the sample spreading module and/or the implementation of time-resolved mixing procedures^24-26^ to trigger (photo-chemical) reactions on grids with a time resolution in the millisecond range^27-29^. This would facilitate capturing and imaging short-lived reaction intermediates with time-resolved stop-motion cryo-EM to elucidate the biochemical mechanisms underpinning biomolecular systems.

Finally, we aim at transferring the EasyGrid technology to other facilities and structural biology laboratories. The modularity of EasyGrid simplifies its implementation as either a full platform or as standalone machines encompassing a subset of functionalities. On its own, EasyGrid Control could help laboratories with limited access to cryo-EM microscopes to screen grids and meet the sample quality criteria required to access high-end cryo-EM imaging facilities. As a full platform, EasyGrid would assist cryo-EM/ET facilities with sample screening and supplement their service portfolio with the ability to jet-vitrify cells and to automatically optimize cellular or biochemical samples.

## Materials and Methods / Experimental details

### Apoferritin preparation parameters

Purified apoferritin solution was prepared at the Protein Expression and Purification Facility in EMBL Heidelberg. Apoferritin was dispensed without dilution in two consecutive steps on holey-carbon R2/2 300 mesh copper grids (Quantifoil) plasma-treated 10 times in EasyGrid right before sample dispensing. First, the grids were coated with a layer of sample by spreading a line of 100 droplets dispensed on the grid with a 2.2 bars pressure wave applied on both sides of the grid for the duration of 30 ms, and then leaving the grid to dry for 10 s at room humidity to clear up the foil holes from coating solution. This coat ensured that the next sequence of dispensing and spreading would produce a thin film of solution. Another line of 100 droplets was then dispensed on coated grids and spread at 20 °C and >92% humidity with a 2.2bars pressure wave for the duration of 30 ms. No delay was added between sample spreading and jet-vitrification of the grids. The grids were then shipped in cold nitrogen vapor to the EMBL Imaging Centre for cryo-EM imaging.

### Apoferritin image acquisition and processing

Data acquisition was carried out using a Titan Krios G4i transmission electron microscope (TEM) operated at an acceleration voltage of 300 kV and equipped with a Falcon4i direct electron detector and a SelectrisX energy filter (Thermo Fisher Scientific) operated in zero loss mode with a filter slit size of 10 eV. A total of 4200 micrographs were acquired at a nominal magnification of 165,000x, resulting in a calibrated pixel size of 0.731 Å. The TEM was operated with a spot size of 5 and each target was exposed for 4 seconds, accumulating a total dose of 40 e^-^/Å^2^ in counting mode in 1135 EER frames. The target defocus range was set between -1.6 and -0.6 μm. Motion correction of EER movies was performed with the CPU implementation of MotionCor2 from Relion4^30^. Initial CTF estimation was performed using CTFFIND4^31^. Particle picking was performed using Gautomatch^32^. Particle extraction and Bayesian polishing were performed with Relion3^30^. All other cryo-EM data processing steps were performed in cryoSPARC^33^ 4.2.1. Conversion of cryoSPARC ‘.cs’ files to Relion ‘.star’ files was performed using UCSF pyem^34^ in combination with in-house bash scripts^35^. All final Homogeneous Refinements done in cryoSPARC^33^ had options “per-particle defocus”, “per-group CTF parameter optimization” and “Ewald Sphere Correction” switched on, resulting in a structure resolved at 1.94Å (FSC threshold 0.143) from 188570 particles. Local resolution estimation was performed using Relion4^30^. Cryo-EM density renderings were generated with ChimeraX^36^.

### Ribosome preparation parameters

*Schizosaccharomyces Pombe* K972 yeast cells grown overnight in YES (Yeast Extract with Supplements, Formedium) medium was used to inoculate 500 mL YES to a density of OD_600nm_= 0.05 and grown overnight at 30

°C while shaken at 225 rpm until they reached an OD_600nm_ of 1.5 to 2. Cells were harvested by centrifugation at 3000 g for 5 min at room temperature and the resulting cell pellet was resuspended in sterile milliQ-H_2_O and transferred to a 50 mL centrifugation tube. Following a sedimentation at 3000 g for 5 min at room temperature, the cells were resuspended in lysis buffer (20 mM Hepes pH 7.4, 100 mM KCl, 5 mM Mg(OAc)_2_ containing a protease inhibitor cocktail tablet from Roche) and sedimented again as above. The resulting pellets were resuspended in 2 mL of lysis buffer per 1 g wet weight and lysed using sterilized glass beads (ø3-5 mm) by four cycles of vortexing at 4 °C for 30 seconds, with 30 seconds of cooling on ice between each cycle. Following a sedimentation as above of the cell debris and glass beads, the supernatant was subjected to a clearing spin at 12,000 g for 10 min at 4 °C. The cleared lysate was loaded onto a 50% (w/V) sucrose cushion containing lysis buffer and subsequently centrifuged at 36,100 rpm (SW55Ti rotor, 158,000 g) for 20 hours at 4 °C. Following centrifugation, the ribosome pellet was carefully resuspended in 150 µL cryo-EM buffer (20 mM Hepes pH 7.4, 60 mM KCl, 5 mM Mg(OAc)_2_) and diluted to a concentration of 250 μg/µL as determined by spectrophotometric analysis (A_260nm_). Freshly prepared ribosome samples were applied to cryo-EM grids as follows:

Quantifoil R2/2 300 mesh copper grids were first plasma-treated 15 times in EasyGrid, then coated on half of their surface with Blue Dextran (BD) (Merck; reference D4772) by dispensing a horizontal line of 100 droplets of 2 mg/mL BD at a vertical offset of +0.2 mm from the central axis of the grid and spreading it with a 2.2 bars pressure wave at >80% humidity and 4 °C for the duration of 30 ms, then leaving the grid to dry at room humidity and temperature for 10 s to clear up the foil holes from coating solution. After coating, pure ribosome solution was applied on the grids as a line of 100 droplets dispensed at an offset of +0.3 mm from the central axis of the grid and spread with a pressure wave for the duration of 50 ms at >80% humidity and 4 °C. No delay was added between sample spreading and jet-vitrification of the grids. The grids were then shipped in cold nitrogen vapor to the EMBL Imaging Centre for cryo-EM imaging.

### Ribosome image acquisition and processing

Data acquisition was carried out using a Titan Krios G4i TEM operated at an acceleration voltage of 300 kV and equipped with a Falcon4i direct electron detector and a Selectris energy filter (Thermo Fisher Scientific) operated in zero loss mode with a filter slit size of 10 eV. A total of 9747 micrographs were acquired at a nominal magnification of 165,000x, resulting in a calibrated pixel size of 0.731 Å. The TEM was operated with a spot size of 4 and each target was exposed for 1.6 seconds, accumulating a total dose of 48 e_-_/Å^2^ in counting mode in 504 EER frames. The target defocus range was set between -1.6 and -0.6 μm. Motion correction of EER movies was performed with the CPU implementation of MotionCor2 from Relion4^30^. Initial CTF estimation was performed using CTFFIND4^31^. Particle picking was performed using in-house python scripts applying bandpass filtering and peak detection to detect ribosomal particles in the micrographs^37^. Particle extraction and 3D classification with alignment were performed with Relion4. All other cryo-EM data processing steps were performed in cryoSPARC^33^ 4.2.1. All final Homogeneous Refinements were done in cryoSPARC^33^ with per-particle defocus and per-group CTF parameter optimization as well as Ewald Sphere Correction switched on, resulting in a structure resolved at 2.43Å (FSC threshold 0.143) from 135,490 particles. Local resolution estimation was performed using Relion4. Cryo-EM density renderings were generated with ChimeraX^36^.

### Rhodopsin preparation parameters

The KR2 rhodopsin was produced as previously described^17^. In brief, KR2-coding DNA was first optimized for *E.coli* codons using GeneArt (Thermo Fisher Scientific). That gene was then synthesized commercially (Eurofins) and subcloned into a pET15b plasmid with a C-terminal 6xHis-tag. *E.coli* cells of C41 strain were transformed with pET15b plasmid containing the *KR2* gene. Transformed cells were grown at 37 °C in shaking baffled flasks in the auto-inducing medium ZYP-5052^38^ containing 10 mg/L ampicillin. Cells were induced at an OD_600_ of 0.6-0.7 with 1 mM isopropyl-β-D-thiogalactopyranoside (IPTG). Subsequently, 10 μM all-*trans*-retinal was added and the incubation continued for 3 hours. The cells were collected by centrifugation at 5000 g for 20 min. Collected cells were disrupted in an M-110P Lab Homogenizer (Microfluidics) at 12,000 p.s.i. in a buffer containing 20 mM Tris-HCl, pH 7.5, 5% glycerol, 0.5% Triton X-100 (Sigma-Aldrich) and 50 mg/L DNase I (Sigma-Aldrich). The membrane fraction of the cell lysate was isolated by ultracentrifugation at 125,000 g for 1 h at 4 °C. The pellet was resuspended in a buffer containing 50 mM Tris-HCl, pH 8.0, 0.2 M NaCl and 1% DDM (Anatrace, Affymetrix) and stirred overnight for solubilization. The insoluble fraction was removed by ultracentrifugation at 125,000 g for 1h at 4 °C. The supernatant was loaded on an Ni-NTA column (Qiagen), and the protein was eluted in a buffer containing 50 mM Tris-HCl, pH 8.0, 0.2 M NaCl, 0.4 M imidazole and 0.1% DDM. The eluate was subjected to size-exclusion chromatography on a Superdex 200 Increase 10/300 GL (Cytiva) in a buffer containing 50 mM Tris-HCl, pH 8.0, 0.2 M NaCl and 0.05% DDM. In the end, the protein was concentrated to 60 mg/mL using 100,000 MWCO Amicon Ultra-15 filters (Millipore) and stored at –80 °C. For the preparation of the cryo-EM grids the protein was diluted to the concentration of 10 mg/mL using the buffer containing 50 mM Tris-HCl, pH 8.0, 0.1 M NaCl and 0.03% DDM.

Quantifoil R2/2 300 mesh copper grids were first plasma-treated 15 times in EasyGrid, then coated twice on half of their surface with Blue Dextran (BD) (Merck; ref. D4772) by dispensing a horizontal line of 100 droplets of 2 mg/mL BD at a vertical offset of +0.2 mm from the central axis of the grid and spreading it with a 2.2 bars pressure wave at >80% humidity and 4 °C for the duration of 30 ms, then leaving the grid to dry at room humidity and temperature for 10 s. Rhodopsin solution at 10 mg/mL was then applied on the grids as a line of 100 droplets dispensed at an offset of +0.3 mm from the central axis of the grid and spread with 80 ms 2.2 bars pressure wave at >90% humidity and 4 °C. A delay of 500 ms was added between sample spreading and jet-vitrification of the grids.

### Rhodopsin image acquisition and processing

Data acquisition was carried out using a Krios G3 TEM (Thermo Fisher Scientific, European Synchrotron Radiation Facility, Grenoble) operated at an acceleration voltage of 300 kV and equipped with a Gatan K3 direct electron detector and a Gatan Quantum energy filter operated in zero loss mode with a filter slit size of 20 eV. A total of 6671 40-frames movies were acquired at a nominal magnification of 105,000x, resulting in a calibrated pixel size of 0.839 Å. The TEM was operated with a spot size of 6 and each micrograph was exposed for 1.65 seconds, accumulating a total dose of 40.3 e^-^/Å^2^ in counting mode. The target defocus range was set between 0.8 and 2.4 μm, in steps of 0.2 μm. Image data processing was entirely performed using the cryoSPARC^33^ software (4.2.1). Patch motion correction was performed on the raw movies, followed by patch CTF estimation. After exposure curation, 4748 micrographs were selected for further processing. Blob particle picking and particle extraction with a 320x320 pixel box size were performed, followed by a round of 2D classification. Five relevant 2D classes exhibiting different particle orientations were selected and used as templates for a subsequent template picking step. 1,916,859 particles were picked and extracted using a 320x320 pixel box size, then binned to 160x160 pixel box size. Two additional rounds of 2D classification and particle selection were then performed, resulting in a homogenous set of 333,078 particles that were then unbinned. A non-symmetrized ab-initio initial model was generated using a subset of 179,700 particles and used to perform homogenous refinement with imposed C5 symmetry, resulting in a structure resolved at 2.36 Å (FSC threshold 0.143). Local CTF Refinement and a final round of Local Refinement were performed, improving the resolution to 2.32 Å (FSC threshold 0.143). Cryo-EM density renderings were generated with ChimeraX^36^.

### Cell culture, vitrification and lamella milling

HeLa Tokyo cells were grown at 37 °C in DMEM medium (Invitrogen) supplemented with penicillin, streptomycin and fetal bovine serum. SUM159 (human breast cancer) cells were maintained in DMEM/F-12 GlutaMAX (Thermo Fisher Scientific) supplemented with 5 mg/mL insulin (Cell Applications), 1 mg/mL hydrocortisone (Sigma), 5% FBS (v/v), 50 mg/mL streptomycin and 50 U/mL penicillin. SiO_2_ R1.2/20 gold grids (Quantifoil) were micro-patterned using a PRIMO device (Alveole) following a previously described protocol^20^. To seed cells on grids, confluent cell cultures were rinsed in PBS and treated with 0.5 mL trypsin for 2 min before seeding them onto grids. For SUM159 cells, 4.0 × 10^5^ trypsinized cells were seeded onto grids in 35 mm low µ-Dishes (ibidi) (4–5 grids per dish) and incubated for 20–30 min. After cell attachment, grids were transferred to another dish with fresh DMEM. The next day, grids were inserted one at a time into the EasyGrid loading station, then immediately picked-up by the gripper robot and transferred to the high-humidity chamber maintained at 30 °C and >75% humidity to limit cell stress induced by DMEM evaporation. The grids prepared with EasyGrid were then treated with a 1.8 bars pressure wave for the duration of 400 ms and immediately jet-vitrified. The total duration between picking a cell-seeded grid from the culture dish and jet-freezing it did not exceed 40 s. The plunge-frozen grids were prepared with a Leica GP2 plunger (Leica Microsystems) maintained at 37 °C and >90% humidity. Grids seeded with SUM159 cells were supplemented with 3 µL of culture medium before blotting. Grids were blotted for 2 s from the reverse side of the grid, then plunged into liquid ethane maintained at -182 °C and stored in LN_2_ until further processing. Grids were screened in EGC and inserted into an Aquilos FIB/SEM (ThermoFisher). Lamellae and stress relief trenches^39^ were automatically milled at mid-height of cells located at the center of grid squares with an angle of 13° using serialFIB^40^, followed by manual polishing down to ≈190 nm thickness.

### In-cell cryo-ET

Tilt series were acquired in 14 lamellae of HeLa cells prepared with EasyGrid, using a Titan Krios G3i transmission electron microscope (ThermoFisher) operated at 300 kV and equipped with a Bioquantum energy filter and a K3 direct electron detector (Gatan) operated in zero-loss mode. On each grid, lamellae were mapped with a pixel size of 28Å, -100 μm defocus, a 70 μm objective aperture and a 30eV energy slit. 14 lamellae were mapped for tilt series (TS) acquisition in SerialEM^41^ following an established protocol^42^. Data was acquired at a nominal magnification of 42,000x resulting in a calibrated pixel size of 2.075Å on the camera. A 50 μm C2 and a 70 μm objective aperture were inserted and the width of the energy filter slit was set to 10 eV. The TEM was operated in nanoprobe mode at spot size 7. TS were acquired using dose-symmetric tilt-scheme with 2° tilt increment and tilt angles ranging from -35° to +61° centered on lamella pretilt (+13°). Movies were acquired in counting mode over 270 ms exposure time with a total accumulated dose of ∼2.63 e-/Å2 per movie (∼15 e-/px/s over an empty area on the camera level) and saved in the TIF file format. 11 frames were saved in each raw tilt image. The accumulated dose of each TS amounted to 130 e-/Å2. The target defocus was set from −2 µm to −4 μm with 0.5 μm steps between TS. Tomograms were aligned and reconstructed using AreTomo^43^ v1.1.1 and visually inspected using IMOD^44^ v4.12.17.

### Ice quality assessment by cryo-EM raster imaging of lamellae

Tokyo HeLa cells were either plunge-frozen with a GP2 plunger (Leica) or prepared with EasyGrid, followed by lamella milling as described above. Lamellae were then imaged using a Titan Krios G4i TEM (ThermoFisher) operated at 300 kV and equipped with a SelectrisX energy filter and a Falcon4i direct electron detector (ThermoFisher) operated in zero-loss mode. A raster of TEM images spaced by 1 µm from one another and avoiding ice-contaminated areas and the platinum layer at the front of the lamella were acquired on each lamella with a pixel size of 1.189Å and a total dose of 20 e^-^/Å^2^. Five images were acquired for each position to overcome possible drift, resulting in 18160 images at 3632 positions on 20 lamellae from 20 plunge-frozen cells from 2 different grids, and 11688 images at 2336 positions on 16 lamellae from 16 jet-vitrified cells from 2 different grids. Frames acquired in .eer format were motion-corrected and converted to .mrc files using Relion4^30^. The power spectra of these frames were computed using the IMOD function ‘clip spectrum’ and analyzed using python scripts^37^. Spectral components around the 3.7Å ring of amorphous ice scattering^21^ were masked and further analyzed. Power peaks were detected using signal thresholding with a fixed threshold for all considered images. Resulting hits were first eroded to limit false positives and then dilated to merge neighboring hits into peaks corresponding to the FFT data. Individual peaks were counted in each spectrum using connected component analysis. A crystallinity score was calculated by summing the peak counts for all five images per position. Positions with a crystallinity score <10 were automatically labeled ‘vitreous’, those >50 were labeled ‘crystalline’, and the others were labeled ‘semi-crystalline’ (Suppl. Fig S8). Each image was then visually checked for the presence or absence of crystalline artifacts in the EM image and peaks in the FFT, and re-labeled by a trained microscopist to limit false attributions. This curation modified [vitreous/semi-crystalline/crystalline] proportions from [15/37/48] to [11/24/65] for plunge-frozen cells, and from [75/14/11] to [77/14/9] for jet-vitrified cells (Fig. 5b).

Lamellae were milled in SUM159 cells following a similar protocol. N=17 lamellae were milled in 6 grids of cells prepared with a GP2 plunger (Leica) and N=18 lamellae were milled in 4 grids of cells prepared with EasyGrid. Qualitative scores from 0 to 5 were attributed to each lamella depending on crystallinity and usability (Suppl. Fig S7).

## Supplementary material

**Fig. S1.**
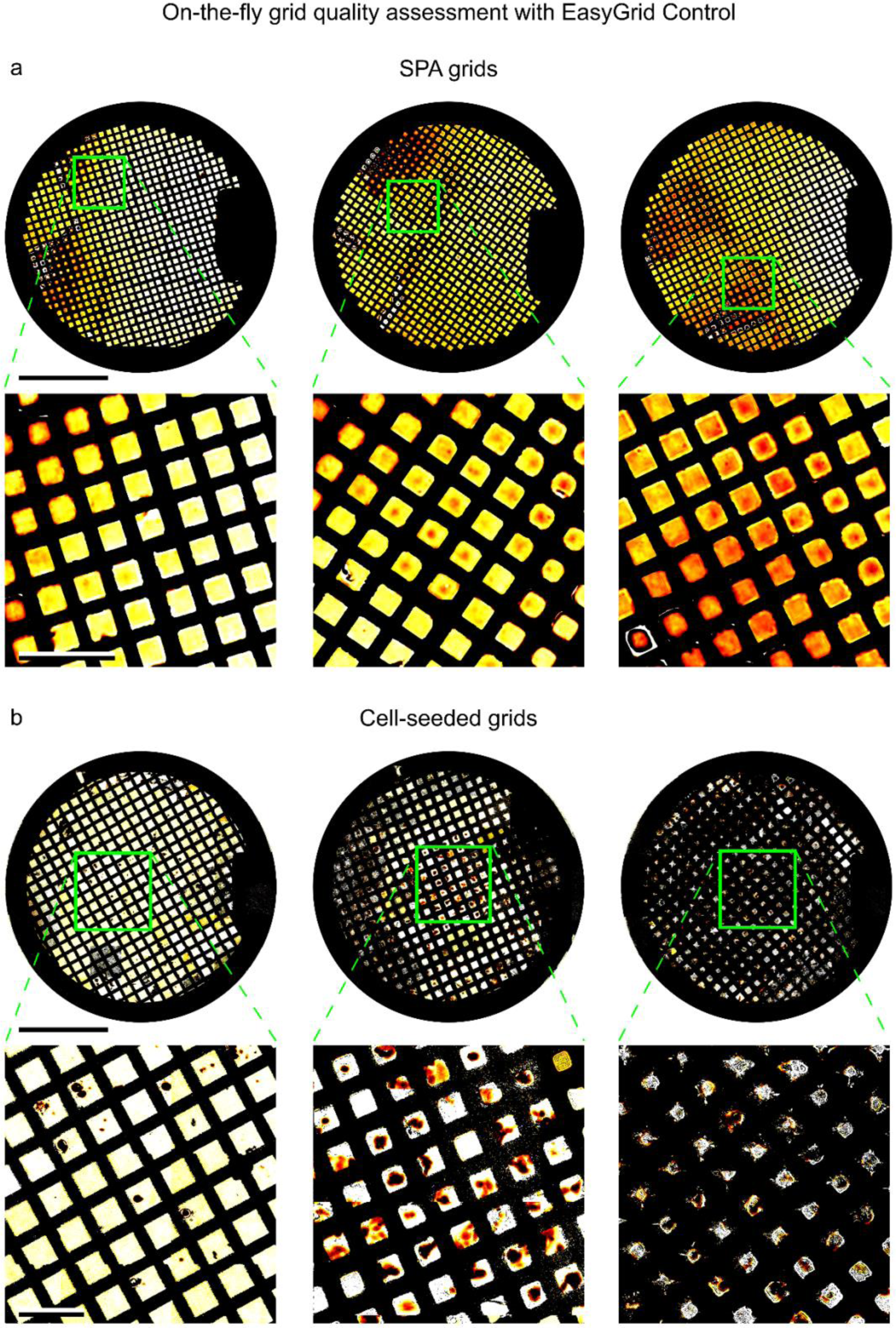
EasyGrid Control enables on-the-fly selection of optimal grids. a. EGC atlases of SPA grids. The gradient of thin ice in the 1^st^ inset suggests a potentially high-quality region for cryo-EM imaging; scale bars, 1 mm (top row) and 200 µm (bottom row). Colormap in arbitrary units (see Discussion). b. EGC atlases of cell-seeded grids. Thicker regions on the 2^nd^ and 3^rd^ atlases indicate that cells remained hydrated throughout sample processing; scale bars, 1 mm (top row) and 200 µm (bottom row).

**Fig. S2.**
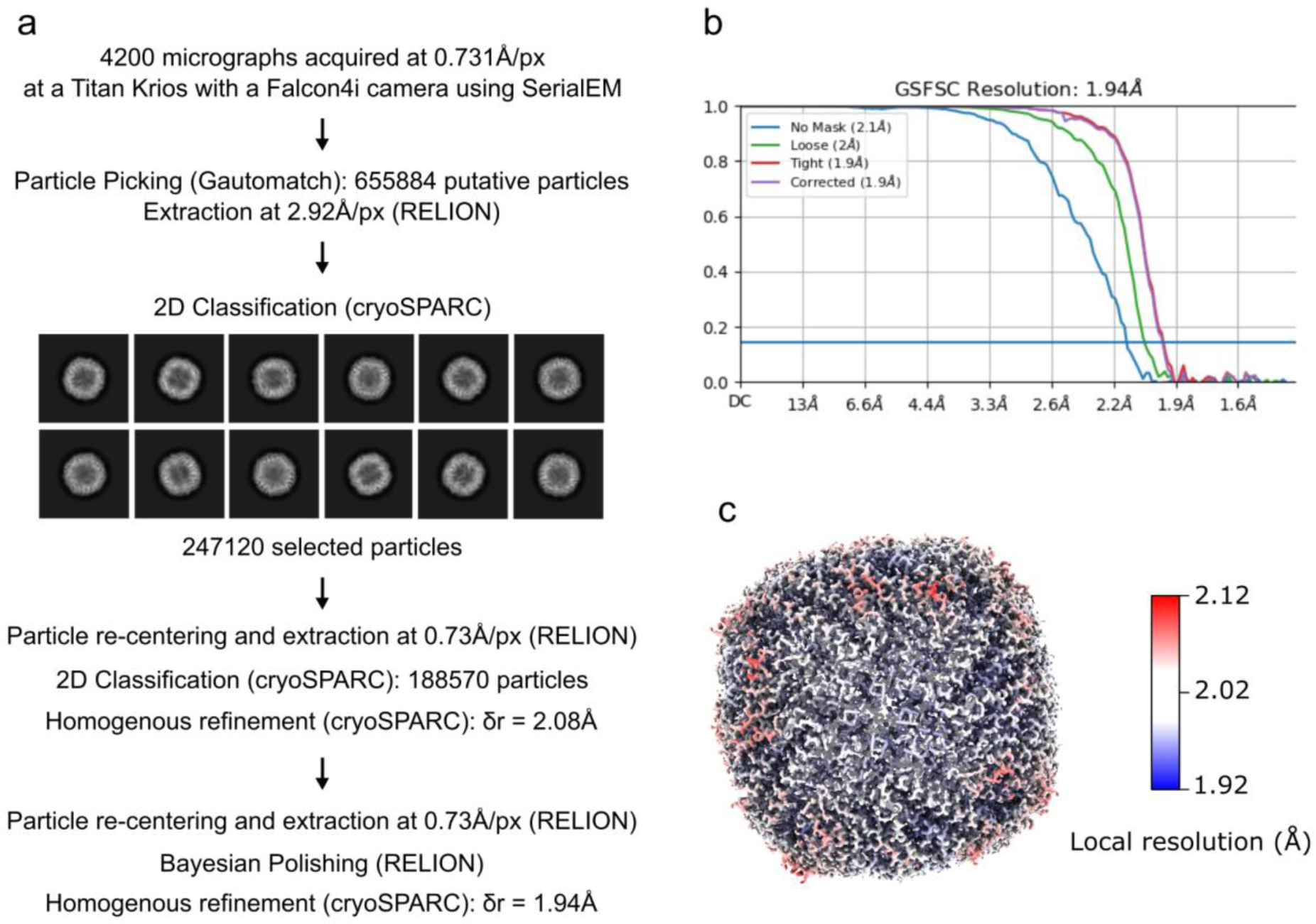
Processing scheme of the single-particle analysis of apoferritin prepared with EasyGrid. a. SPA processing workflow for the apoferritin data. 656k particles were picked from motion-corrected and dose-weighted micrographs using Gautomatch and extracted from micrographs binned 4 times (pixel size of 2.92Å) using RELION. A subset of 247k particles were selected using 2D-classification in cryoSPARC, then re-extracted from unbinned micrographs and submitted to a second round of 2D-classification. The selected 189k particles underwent Homogenous Refinement in cryoSPARC. Re-centered particle stacks were extracted in RELION and underwent Bayesian polishing in RELION, then Homogenous Refinement in cryoSPARC, yielding a cryo-EM density map of apoferritin at 1.94Å resolution (δr). b. Gold-standard Fourier shell correlation (GSFSC) plot for the final half-maps. The horizontal blue line indicates the resolution estimation threshold at 0.143. c. Final cryo-EM density map of apoferritin color-coded by local resolution computed in RELION and displayed in ChimeraX.

**Fig. S3.**
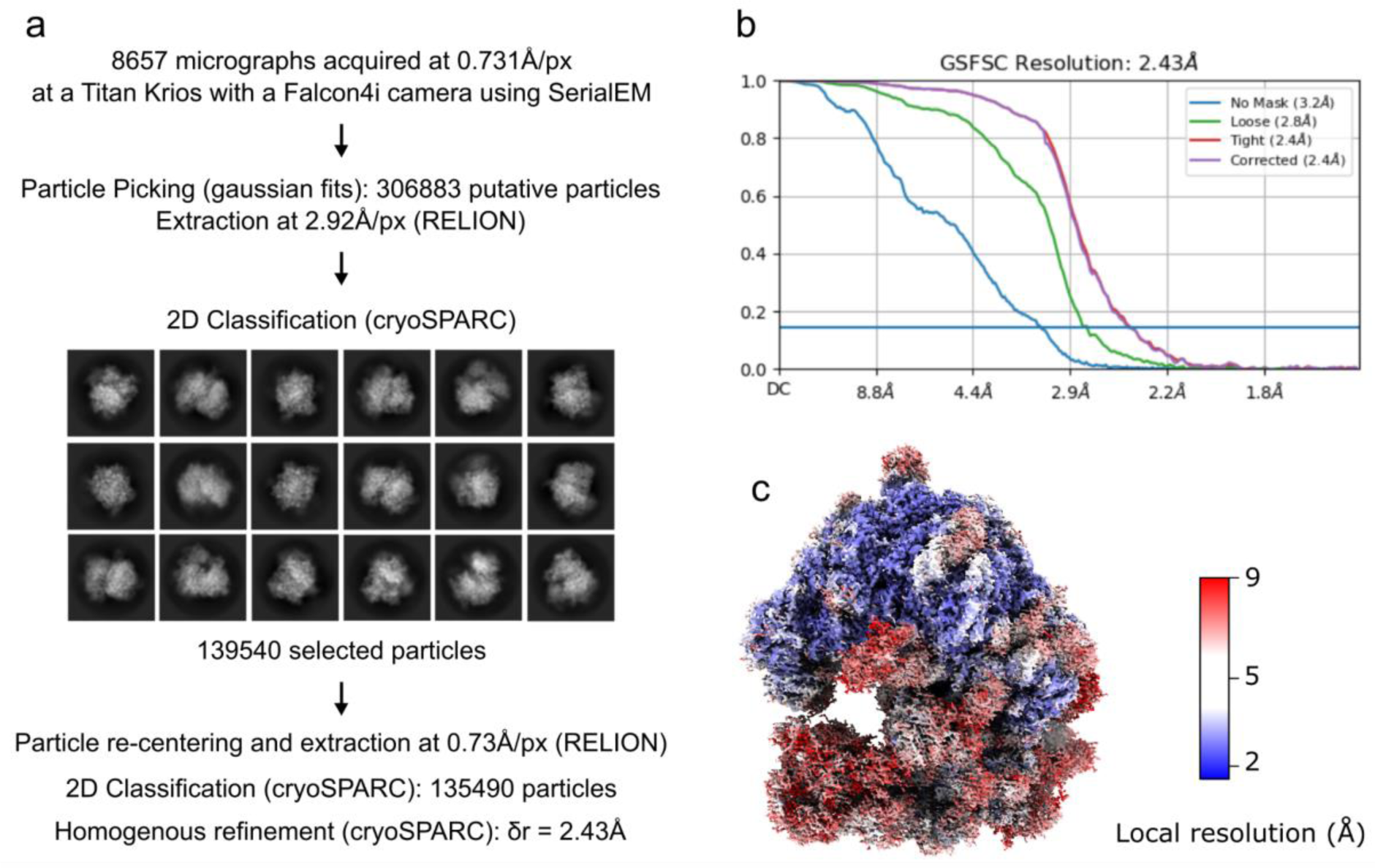
Processing scheme of the single-particle analysis of yeast ribosome prepared with EasyGrid. a. SPA processing workflow for the ribosome data. 307k particles were picked from motion-corrected and dose-weighted micrographs using python-based picking of signal densities measuring ∼300Å, then extracted from micrographs binned 4 times (pixel size of 2.92Å) using RELION. A subset of 140k particles were selected using 2D-classification in cryoSPARC, then re-extracted from unbinned micrographs and submitted to a second round of 2D-classification. The selected 135k particles underwent Homogenous Refinement in cryoSPARC, yielding a cryo-EM density map of the full ribosome at 2.43Å resolution (δr). b. Gold-standard Fourier shell correlation (GSFSC) plot for the final half-maps. The horizontal blue line indicates the resolution estimation threshold at 0.143. c. Final cryo-EM density map of the ribosome color-coded according to local resolution computed in RELION and displayed in ChimeraX. Because the small ribosomal subunit (SSU) can ratchet relative to the large ribosomal subunit^45^ (LSU), and particle alignment was driven by the LSU, the resolution of the SSU is comparatively lower (white/red domain at the base of the map).

**Fig. S4.**
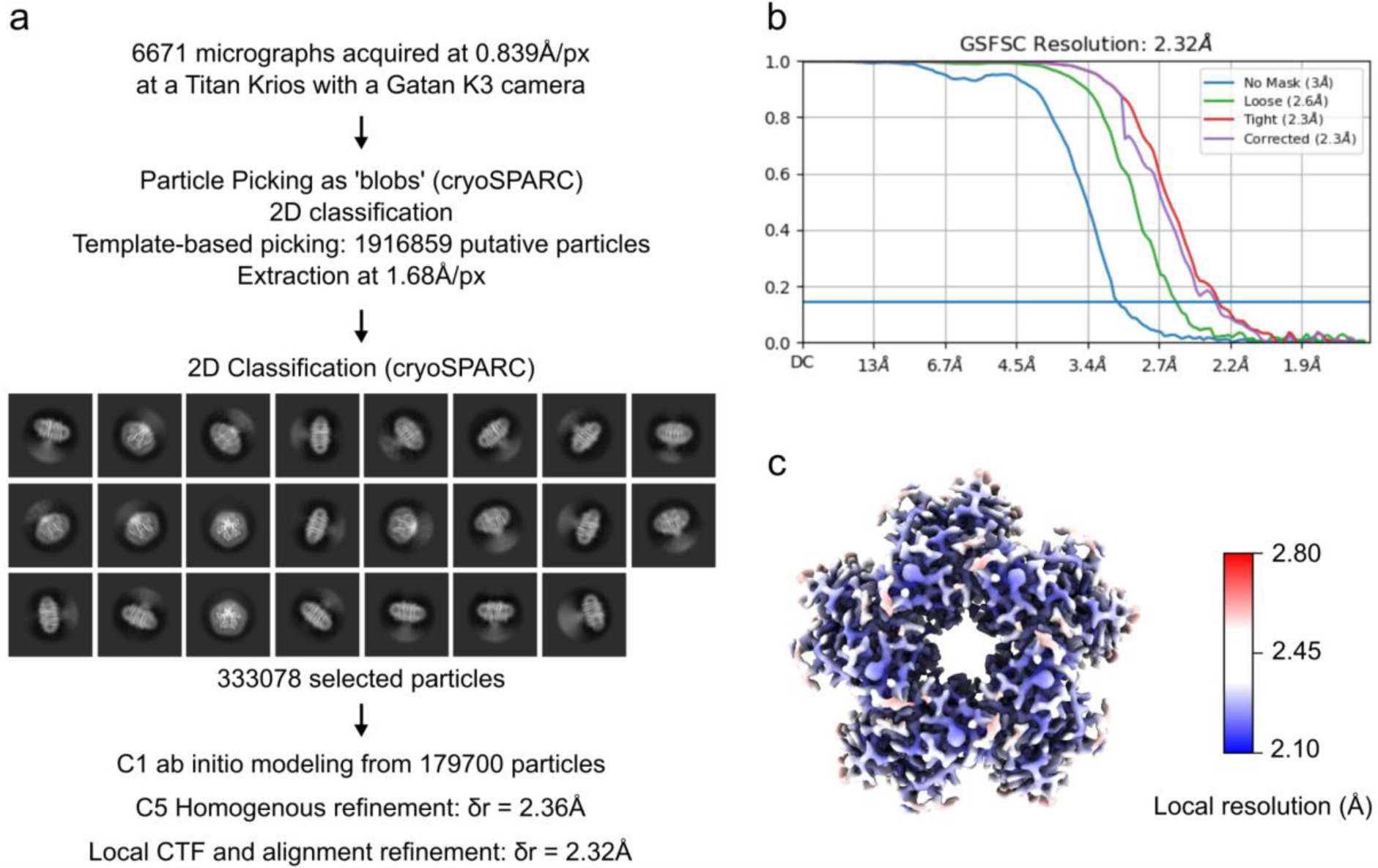
Processing scheme of the single-particle analysis of rhodopsin prepared with EasyGrid. a. SPA processing workflow for the KR2 rhodopsin data, performed exclusively with cryoSPARC. 1917k particles were picked from motion-corrected and dose-weighted micrographs, then extracted from micrographs binned 2 times (pixel size of 1.68Å). A subset of 333k particles were selected using 2D-classification. 180k particles from those were used to generate a non-symmetrized (C1) *ab initio* model of the rhodopsin pentamer, which allowed structure refinement with C5 symmetry for the 333k particles. Subsequent local refinement yielded a cryo-EM density map of the KR2 rhodopsin pentamer at 2.32Å resolution (δr). b. Gold-standard Fourier shell correlation (GSFSC) plot for the final half-maps. The horizontal blue line indicates the resolution estimation threshold at 0.143. c. Final cryo-EM density map of the rhodopsin complex color-coded by local resolution computed in cryoSPARC and displayed in ChimeraX.

**Fig. S5.**
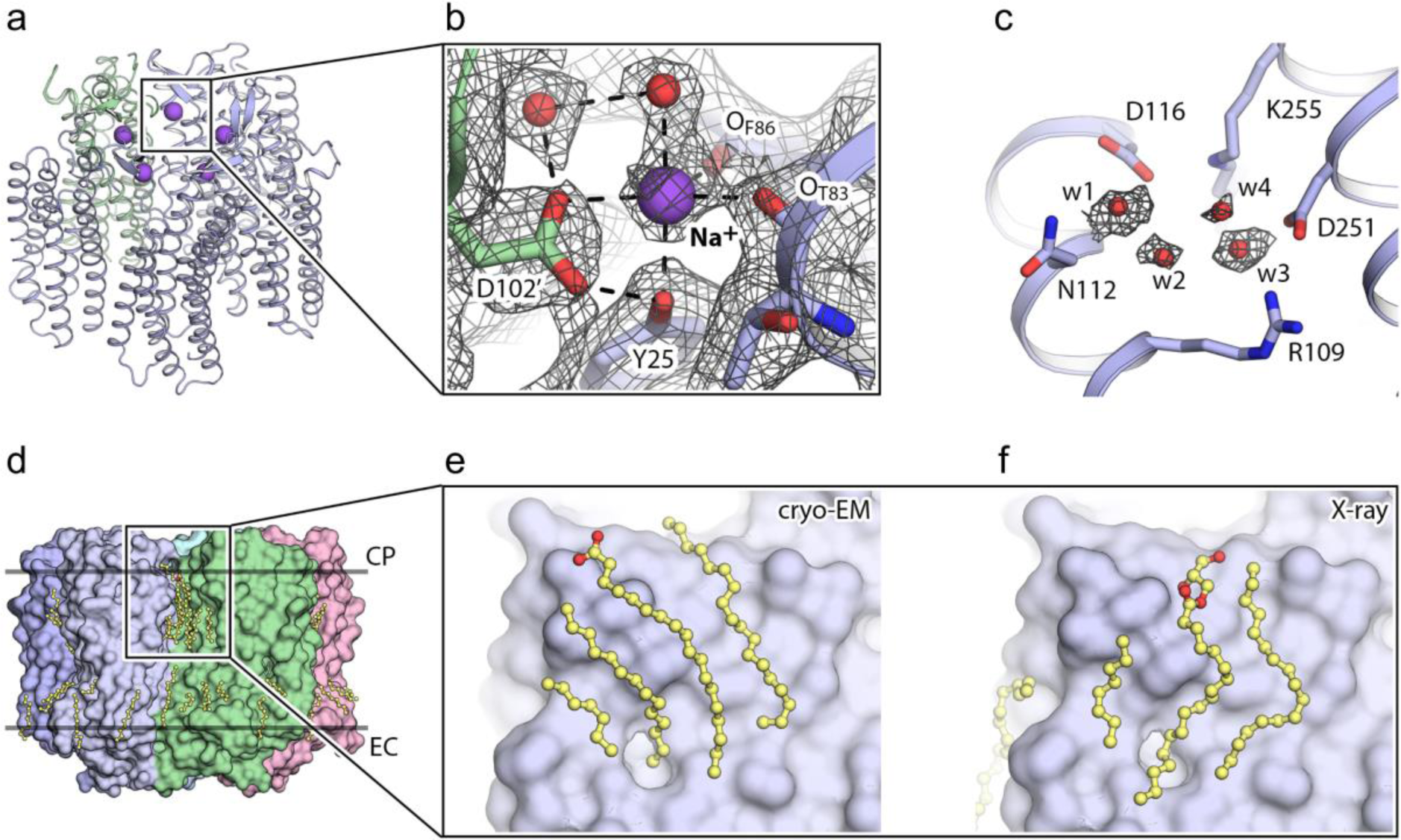
Cryo-EM reveals the high variability of lipid conformation on KR2 rhodopsin complexes. a. Overview of the KR2 pentamer with bound interprotomeric sodium ions shown as purple spheres. b. Detailed view of the interprotomeric sodium binding site. Cryo-EM densities were contoured at the level of 4 root mean square deviations (r.m.s.d.) and displayed as a black mesh. c. View of the active center of KR2. N112 is in the expanded conformation. The cryo-EM densities depicting the 4 water molecules in the internal cavity are shown as a black mesh and contoured at 4 r.m.s.d. d. Side view of the KR2 pentamer featuring lipidic molecules (yellow sticks and spheres) visible in the cryo-EM map. CP: cytoplasmic side; EC: extracellular side. e. Close-up on lipid fragments located on the cytoplasmic half of the interprotomeric cleft, as revealed by our cryo-EM study. f. Close-up on lipid fragments located on the cytoplasmic half of the interprotomeric cleft, as described in a previous study using X-ray crystallography (PDB ID: 6YC3).

**Fig. S6.**
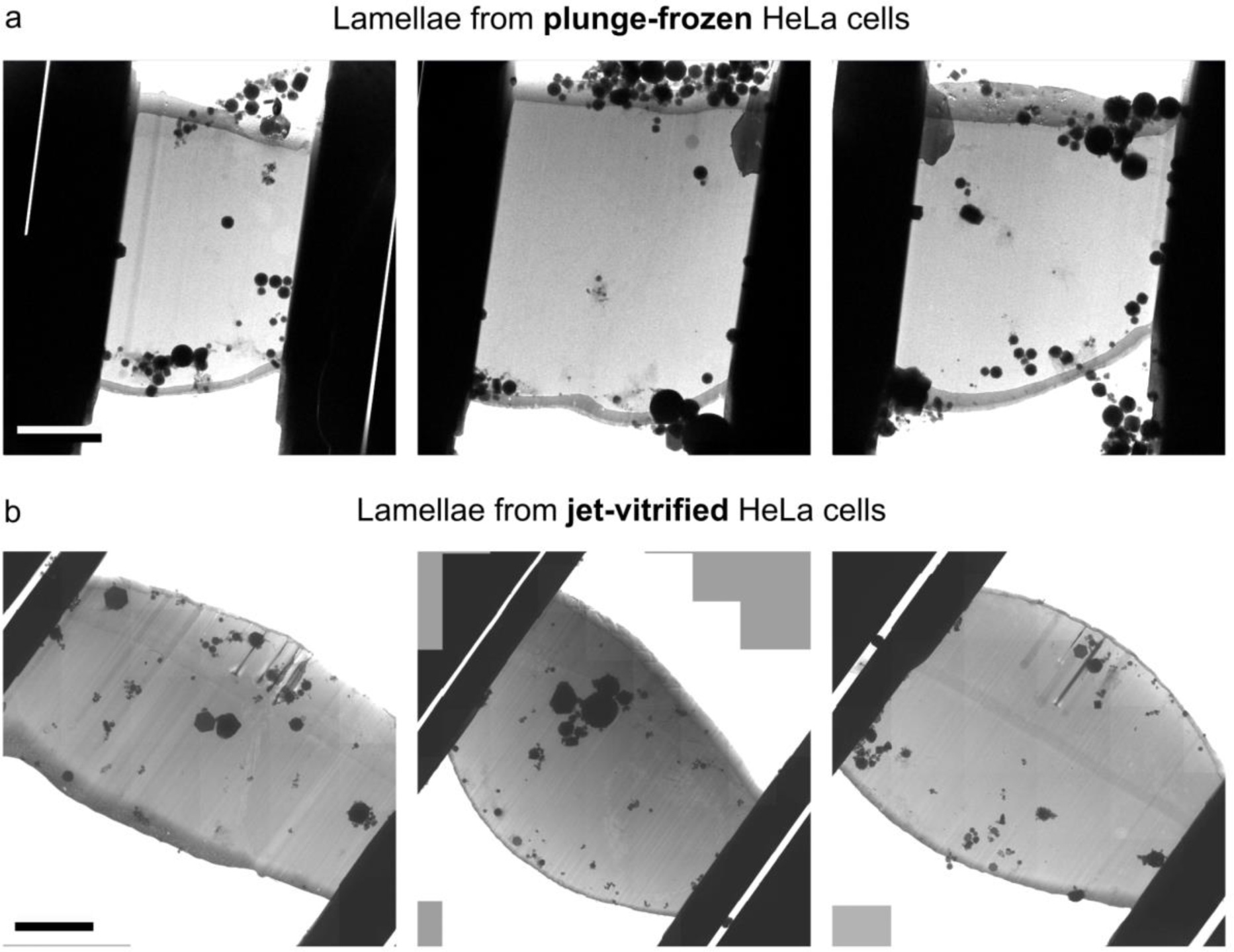
Examples of lamellae milled in plunge- and jet-vitrified HeLa cells. a. TEM images of lamellae from plunge-frozen HeLa cells prepared with Leica GP2; scale bar, 5 µm. b. TEM images of lamellae from jet-vitrified HeLa cells prepared with EasyGrid. Quantification of ice quality based on raster-imaged at 1.2Å/px is provided in Figure 5.

**Fig. S7.**
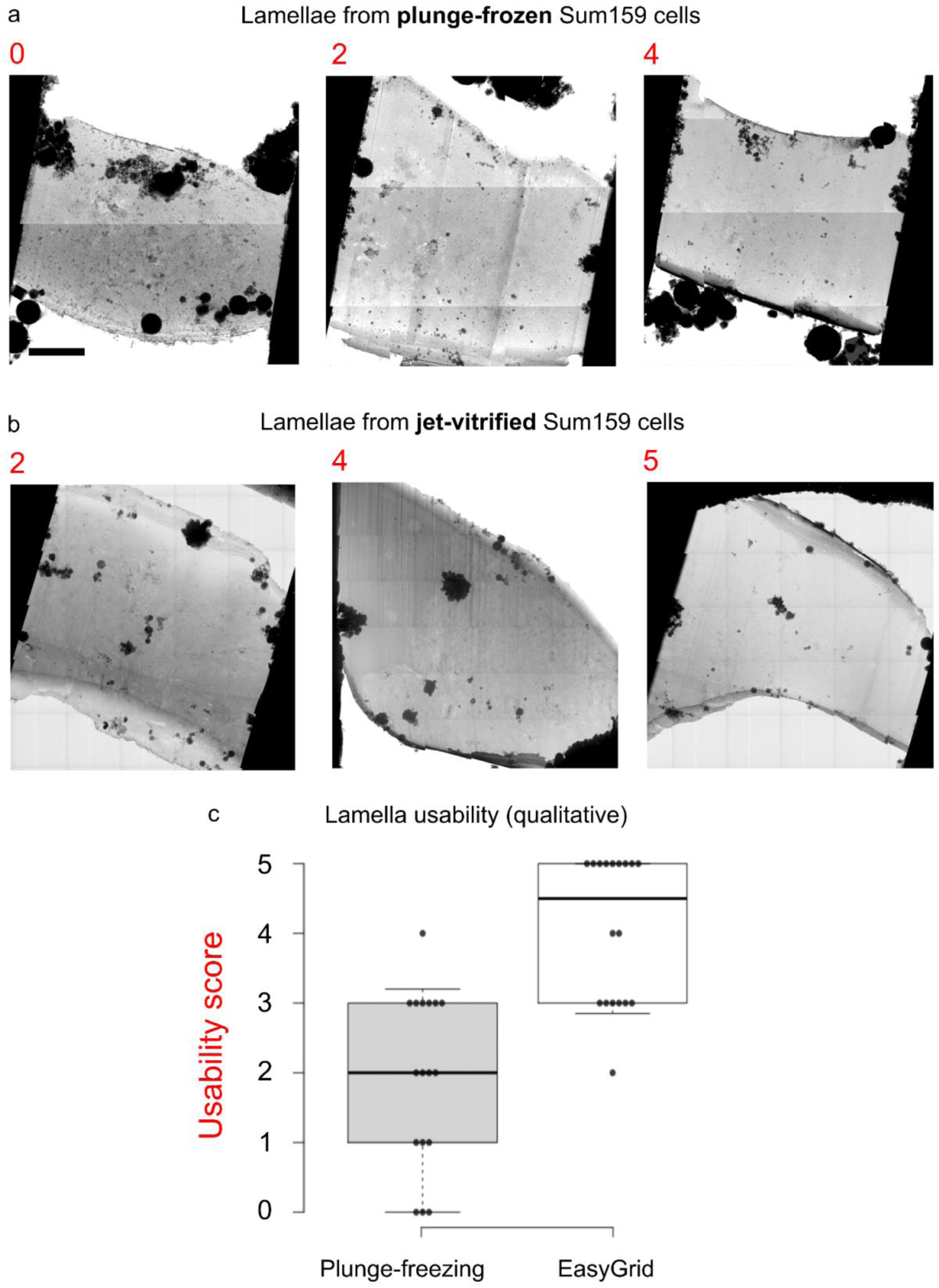
Qualitative assessment of ice quality in plunge- and jet-vitrified SUM159 cells. a. TEM images of lamellae from plunge-frozen SUM159 cells prepared with Leica GP2. A qualitative score (red number) was attributed to each lamella based on visual inspection. The score ranges from 0 describing a completely crystalline, unusable lamella to 5 describing a vitreous, high-quality lamella – independently of milling artifacts; scale bar, 5 µm. b. TEM images of lamellae from jet-vitrified SUM159 cells prepared with EasyGrid. c. Qualitative comparison of lamella usability in lamellae prepared by either plunge- or jet-vitrified SUM159 cells; n=17 and 18 lamellae respectively.

**Fig. S8.**
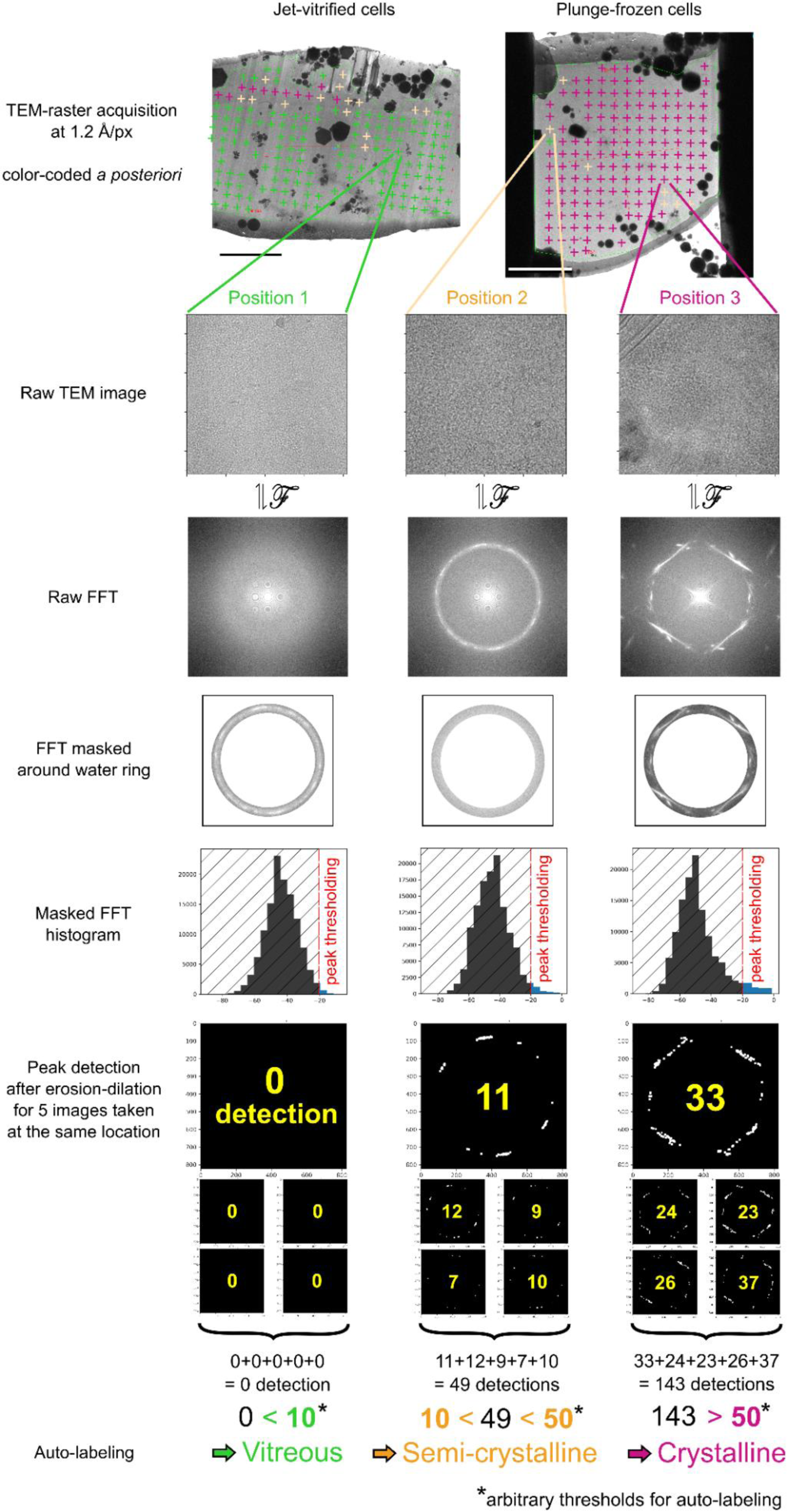
Ice quality quantitation analysis scheme. Step-by-step analysis workflow of ice crystallinity in lamellae using raster-TEM imaging, exemplified with images taken at three different positions and exhibiting either vitreous (left column), crystalline (right column) or ambiguous signal (middle column). At each location marked by a cross on the left-most panels, 5 TEM images were acquired at 1.2Å pixel size to account for the observed variation of the 2D Fourier transforms upon repeated acquisition. In the Fourier power spectrum of those images, the water ring was masked and diffraction peaks were detected in this region by thresholding out low power values, eroding away single pixels to remove shot noise, dilating peaks back to their original size in the diffraction signal, and performing connected component analysis to count the number of diffraction peaks per image. The diffraction score of each sampled position was defined as the total number of peaks detected on all 5 images taken at this location. Empirical thresholds were arbitrarily defined to automatically label positions in three categories: “unambiguously vitreous” (green color, score < 10), “unambiguously crystalline” (magenta color, score > 50) and “ambiguous: thick and/or semi-crystalline” (orange color, all other positions). All images were then visually inspected to correct false attributions, yielding the ice quality quantitation displayed in Fig.5; scale bars, 5 µm.

**Movie S1. EasyGrid live demonstration**

This movie depicts the main functions of EasyGrid for sample preparation with live close-up sequences.

## Data availability

The rhodopsin structure and model were deposited in the Protein Data Bank (PDB) (accession number: 8RQ5). Other maps will be deposited in the Electron Microscopy Data Bank (EMDB) (accession number: XXX). Raw data used for determining the structure of rhodopsin complexes will be deposited in the EMPIAR database (accession number: XXX). Raw data used for ice quality quantification will be deposited in the EMPIAR database (accession number: XXX).

## Code availability

Source code for the ice quality quantification is available on GitHub at https://git.embl.de/gemin/easygrid.

## Supporting information

video of the EasyGrid machine

## Acknowledgements

We thank Michael Adams and Jiangfeng Zhao for providing large amounts of test sample solutions and support with cryo-EM imaging; Arthur Felisaz and Thibault Deckers for EGC image processing; Moritz ‘neverman’ Niemann for his help in preparing the ribosome sample; the EMBL Heidelberg lab kitchen for support; Martin Pelosse and Angélique Fraudeau for support with cell culture. The apoferritin plasmid was a kind gift from Haruaki Yanagisawa (University of Tokyo). The apoferritin solution was prepared by Arne Börgel from the EMBL Protein Expression and Purification Core Facility. We acknowledge continuous support and feedback from the EM Facility in EMBL Grenoble. We thank the EMBL central IT services and Thomas Hoffmann for computational support. We acknowledge the access and services provided by the Imaging Centre at the European Molecular Biology Laboratory (EMBL IC), generously supported by the Boehringer Ingelheim Foundation. We warmly thank Stephen Cusack and Kristina Djinovic Carugo for their continuous support.

## Author contributions

F.C. and G.P. developed the EasyGrid concept with input from M.H., W.G., J.M., C.W.M., S.E. and S.M.

V.A., C.R., K.L., R,J., F.F., J.S., F.C. and G.P. designed, built, implemented and improved the EasyGrid platform.

V.A., O.G., C.B., M.W.B, A.B., L.L. and G.P. prepared grids and helped refine the EasyGrid preparation process.

V.A., T.D., K.L. and G.P. implemented image analysis in the EasyGrid Control module.

O.G., M.H., S.S., R.L., K.K. and L.L. performed cryo-EM imaging and analysis.

O.G., A.B., V.T.S., G.W. prepared FIB-milled cells and/or performed cryo-ET imaging and analysis.

O.G. designed and performed the ice quality quantitation experiment and analysis.

O.G. produced the figures with input from all authors.

O.G., G.P., K.K. and R.L. wrote the manuscript with input from all authors.

